# Re-annotating the EPICv2 manifest with genes, intragenic features, and regulatory elements

**DOI:** 10.1101/2025.03.12.642895

**Authors:** Bethan Mallabar-Rimmer, Jonathan Mill, Eilis Hannon, Amy P Webster

## Abstract

**Motivation:** The Illumina Infinium MethylationEPIC v2.0 BeadChip (EPICv2 array) is a microarray for quantification of DNA methylation at sites across the human genome, succeeding previous iterations of the platform, including the HumanMethylationEPIC BeadChip (EPICv1 array). An open source manifest file provided by the manufacturer maps array probes to genes and regulatory features. However, due to a change in strategy, it is no longer consistent with annotations for previous versions of the technology. We therefore generated an extended EPICv2 manifest, to improve backwards-compatibility with EPICv1 and provide a more comprehensive framework for interpreting DNA methylation data.

**Results:** Using public databases and the genomic coordinates of probes, we mapped the 923,452 sites assayed on the Illumina EPICv2 array, comprehensively annotating genes and regulatory elements. We also replicated the manufacturer’s approach of annotating sites in the regions <=200bp and 201-1500bp upstream of a transcription start site (the TSS200 and TSS1500), ensuring backwards-compatibility with existing pipelines for Illumina methylation array data. We found that 731,759 EPICv2 array sites (79.24% of all sites on the array) are located within a gene body (exon, intron, or UTR) according to the GENCODE Human release 49 (GENCODEv49) database.

We additionally labelled sites located in a promoter or enhancer according to the GeneHancer database. Finally, the re-annotated manifest labels which sites are required for the Horvath DNA Methylation Age Calculator and MethylDetectR epigenetic clocks, to facilitate data preparation for these tools.

**Availability and Implementation:** The re-annotated manifest is freely available at https://doi.org/10.5281/zenodo.14933468. The re-annotation code is on GitHub: https://github.com/bethan-mallabar-rimmer/EPICv2_manifest.

## Introduction

The Illumina Infinium MethylationEPIC v2.0 BeadChip (EPICv2 array), released in 2023, is a microarray developed for genome-wide quantification of DNA methylation. (1) Previous versions of the array (including HumanMethylation450k and EPICv1) rank among the most widely-used technologies for studies of DNA methylation in humans. (2) The EPICv2 array includes 937,690 probes, profiling DNA methylation at 923,452 unique genomic sites, an expansion on the previous version of the technology, the EPICv1 array that included 866,554 probes. (3)

The EPICv2 manifest is an open-source file provided by Illumina, which annotates each probe on the EPICv2 array with technical details about the probe (such as its sequence and Infinium chemistry type), as well as information about the genomic features overlapping the probe’s target site (such as whether the site is located within a gene, promoter, and/or transcription start site, TSS). (4,5) This genomic feature annotation is crucial to the interpretation of all results from studies conducted using EPICv2. A major change between the EPICv2 manifest and manifests for previous Illumina methylation arrays is that sites within introns are no longer annotated with the gene they are located in. Furthermore, these sites are no longer labelled as intragenic.

A change in annotation strategy can introduce issues for studies that look to include data from different array versions for the purposes of meta-analysis, replication, or longitudinal analysis, especially where this is done at the gene level or integration with other ‘omics is required. Therefore, to ensure consistency with previous studies in how probes are annotated to genes, we used GENCODEv49 data (6) to annotate which sites in the manifest are located within genes, transcripts, intragenic features (including introns, exons, coding sequences or CDSs, and untranslated regions or UTRs), or upstream of a transcription start site (TSS). We also label whether sites are located within any predicted enhancer or promoter element according to the GeneHancer database, (7) and indicate which sites are required for upload to the Horvath (8,9) and MethylDetectR (10,11) epigenetic clocks. In summary, we present an updated EPICv2 manifest file that includes comprehensive annotation of intragenic and regulatory features targeted by probes on the array, to improve comparability of EPICv2 and previous methylation arrays and address incomplete coverage in the original manifest.

## Results

### Format of the re-annotated manifest

The re-annotated manifest includes 19 new columns, described in Table 1. To avoid confusion and including redundant information, we have removed existing columns from the manifest containing gene annotations provided by the manufacturer using their new strategy. These columns include: “UCSC_RefGene_Group”, “UCSC_RefGene_Name”, “UCSC_RefGene_Accession”, “GencodeV41_Group”, “GencodeV41_Name”, and “GencodeV41_Accession”. However, a version of the manifest including both our annotation and the manufacturer’s annotation is available to allow comparison between the two, from https://doi.org/10.5281/zenodo.14933468.

**Table 1:**
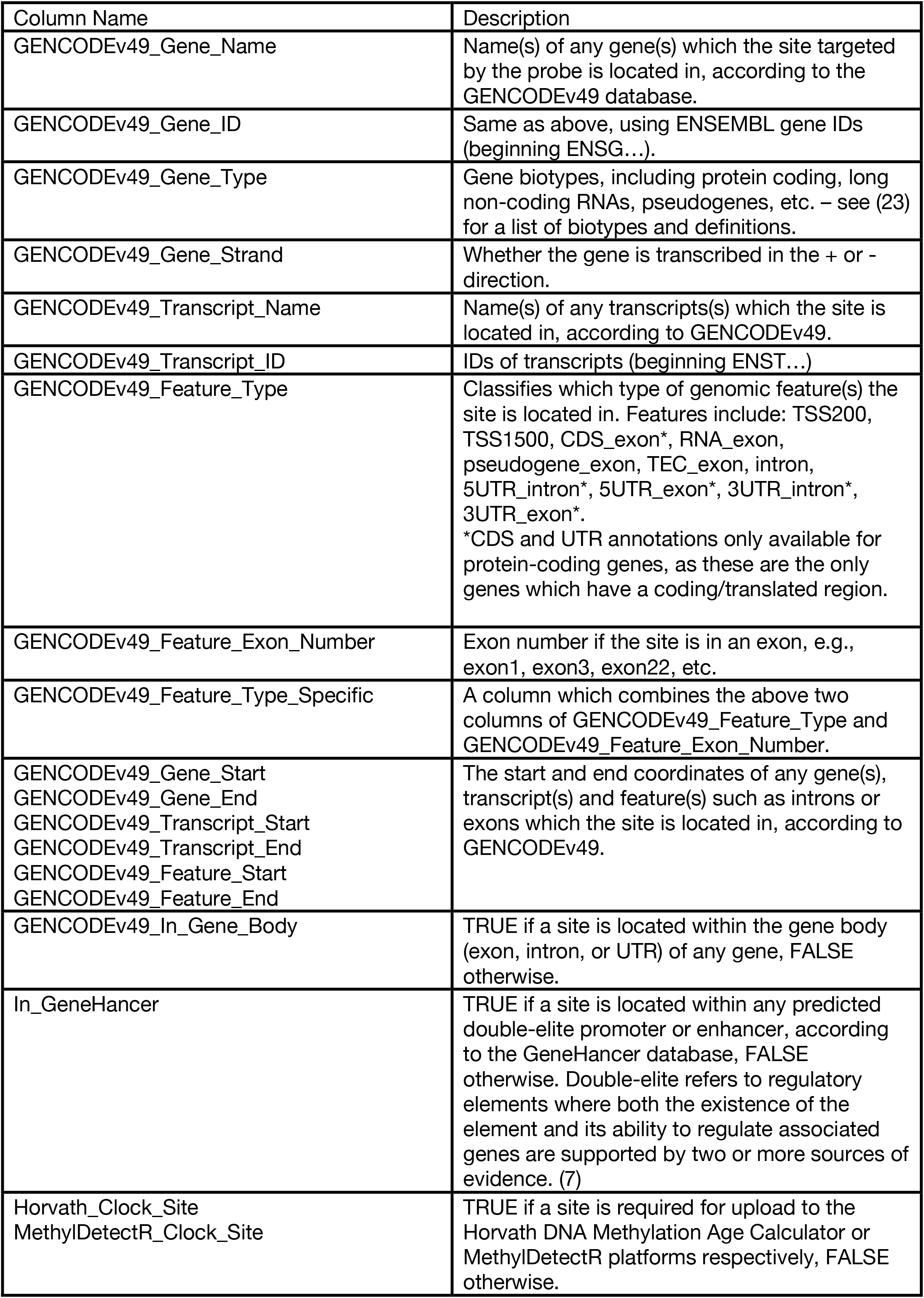
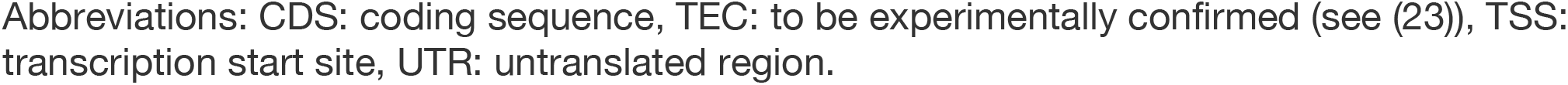
Columns added to the re-annotated EPICv2 manifest, and description of the information provided by each column. See Methods for further detail on the source of these annotations. Abbreviations: CDS: coding sequence, TEC: to be experimentally confirmed (see (23)), TSS: transcription start site, UTR: untranslated region.

| Column Name | Description |
| --- | --- |
| GENCODEv49_Gene_Name | Name(s) of any gene(s) which the site targeted by the probe is located in, according to the GENCODEv49 database. |
| GENCODEv49_Gene_ID | Same as above, using ENSEMBL gene IDs (beginning ENSG...). |
| GENCODEv49_Gene_Type | Gene biotypes, including protein coding, long non-coding RNAs, pseudogenes, etc. – see (23) for a list of biotypes and definitions. |
| GENCODEv49_Gene_Strand | Whether the gene is transcribed in the + or - direction. |
| GENCODEv49_Transcript_Name | Name(s) of any transcripts(s) which the site is located in, according to GENCODEv49. |
| GENCODEv49_Transcript_ID | IDs of transcripts (beginning ENST...) |
| GENCODEv49_Feature_Type | Classifies which type of genomic feature(s) the site is located in. Features include: TSS200, TSS1500, CDS_exon*, RNA_exon, pseudogene_exon, TEC_exon, intron, 5UTR_intron*, 5UTR_exon*, 3UTR_intron*, 3UTR_exon*.<br>*CDS and UTR annotations only available for protein-coding genes, as these are the only genes which have a coding/translated region. |
| GENCODEv49_Feature_Exon_Number | Exon number if the site is in an exon, e.g., exon1, exon3, exon22, etc. |
| GENCODEv49_Feature_Type_Specific | A column which combines the above two columns of GENCODEv49_Feature_Type and GENCODEv49_Feature_Exon_Number. |
| GENCODEv49_Gene_Start<br>GENCODEv49_Gene_End<br>GENCODEv49_Transcript_Start<br>GENCODEv49_Transcript_End<br>GENCODEv49_Feature_Start<br>GENCODEv49_Feature_End | The start and end coordinates of any gene(s), transcript(s) and feature(s) such as introns or exons which the site is located in, according to GENCODEv49. |
| GENCODEv49_In_Gene_Body | TRUE if a site is located within the gene body (exon, intron, or UTR) of any gene, FALSE otherwise. |
| In_GeneHancer | TRUE if a site is located within any predicted double-elite promoter or enhancer, according to the GeneHancer database, FALSE otherwise. Double-elite refers to regulatory elements where both the existence of the element and its ability to regulate associated genes are supported by two or more sources of evidence. (7) |
| Horvath_Clock_Site<br>MethylDetectR_Clock_Site | TRUE if a site is required for upload to the Horvath DNA Methylation Age Calculator or MethylDetectR platforms respectively, FALSE otherwise. |

EPICv2 is the first Infinium methylation array to indicate which probes are replicates, targeting the same genomic site, through its probe naming system. (2,12) Our genomic feature annotation is identical across replicate probes which target the same site.

### Intragenic and regulatory feature annotation

We used GENCODEv49 data to label 737,445 EPICv2 probes (targeting 731,759 sites) as located within a gene body, defined in this analysis as between transcription start and end sites. This annotation encompasses exons, introns, CDSs, and UTRs in protein-coding genes, RNAs, pseudogenes, and “to be experimentally confirmed” (TEC) genes predicted to be protein-coding. Annotation of intraexonic features (CDSs and UTRs) is only provided for protein-coding genes, as only translated genes have a coding and untranslated region.

Using GENCODEv49, 731,759 sites (79.24% of all sites on the array) mapped to the gene body of at least one gene. Of these, 592,721 sites (64.19%) were mapped to a protein-coding gene, 297,903 (32.26%) to an RNA gene, 14,309 (1.55%) to a pseudogene and 3,039 (0.33%) to a TEC gene. 191,780 sites (20.77%) were intergenic i.e. not located within the gene body of any transcript (Figure 1A). 128,789 sites (13.95% of array sites) mapped to the TSS200 and 241,715 (26.18%) mapped to the TSS1500 of at least one transcript. 252,991 (27.40%) mapped <=1500bp upstream (5’) of a protein-coding gene, and 166,986 (18.08%) mapped upstream of an RNA gene (Figure 1B).

**Figure 1.**
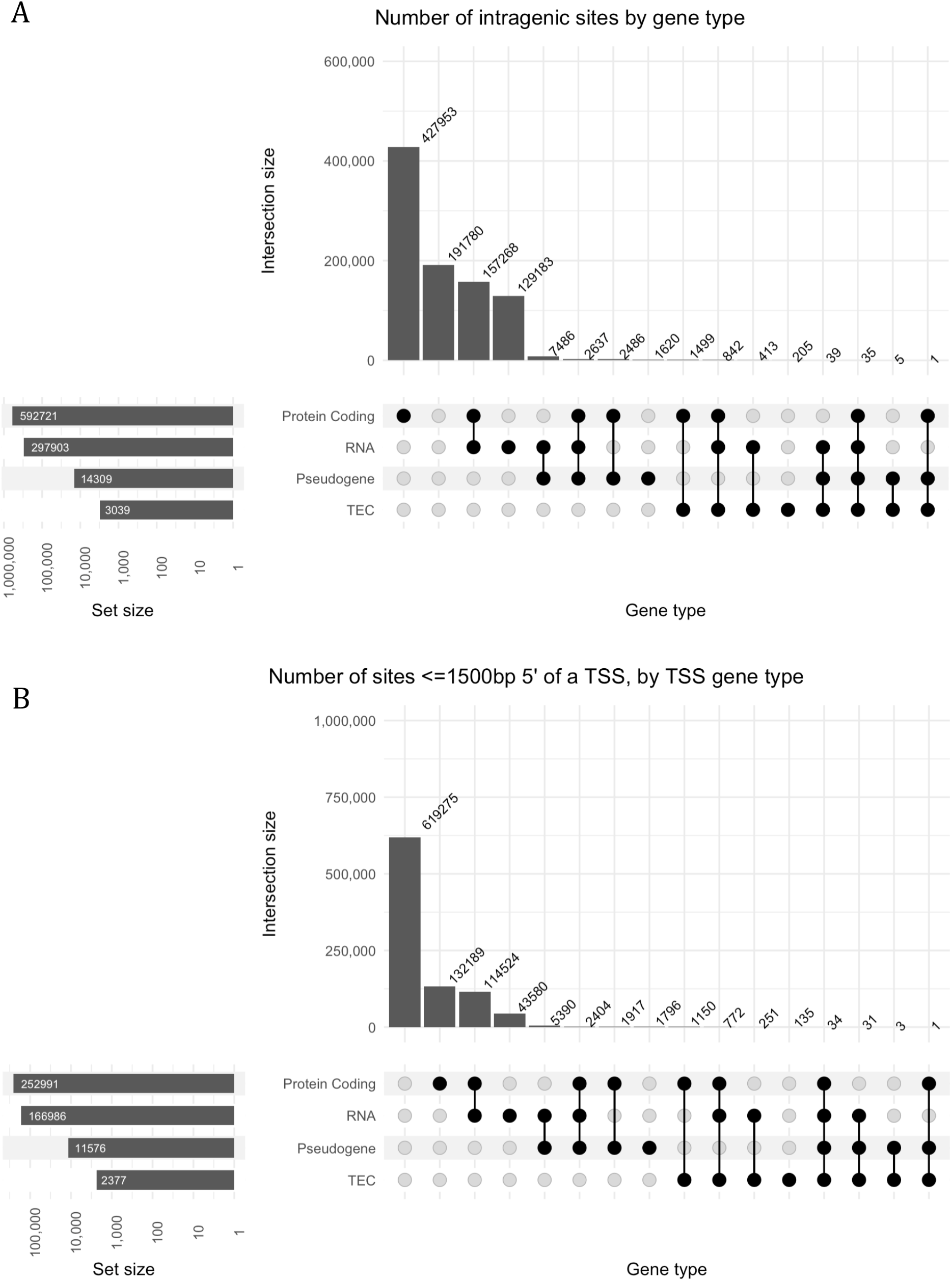
**A:** UpSet plot showing the number of intragenic sites by gene type in the re-annotated manifest. Intragenic refers to sites within a gene body, including those located in an intron or coding/UTR exon. Intragenic sites were most commonly mapped to at least one protein-coding gene and no other gene type (n=427,953), followed by at least one protein-coding and RNA gene (n=157,268). 191,780 sites were not mapped to the gene body of any transcript. For both panels, the “set size” bar plot shows the total number of sites mapped to each gene type. **B:** Numbers of sites in either the TSS200 or TSS1500, by gene type the region is 5’ of, in the re-annotated manifest. The majority of sites were not mapped to the TSS200 or TSS1500 region of any transcript (n=619,275). Of sites mapped <1500bp 5’ of a TSS, most were mapped upstream of at least one protein-coding gene (n=132,189), followed by at least one protein-coding and RNA gene (n=114,524).

We also annotated the regulatory genome using the GeneHancer database, which integrates the Ensembl, Encyclopedia of DNA Elements (ENCODE), Functional Annotation of Mammalian Genomes (FANTOM) and Vista Enhancer Browser databases of promoters and enhancers. (7) Unfortunately, constraints on GeneHancer data sharing mean we are unable to include specific IDs of promoters/enhancers and the genes regulated by these elements in the manifest. (7,13) Instead, our annotation uses a binary true/false measure to indicate whether a site is present in any regulatory element according to GeneHancer. Users can find information about the specific regulatory element and associated genes by viewing the CpG site in the UCSC Genome Browser with the GeneHancer track enabled (14) and information about each regulatory element can found on the GeneCards website. (15,16) Only double elite regulatory elements, with both the existence of the element and its ability to regulate associated genes supported by two or more sources of evidence, are included in our annotation. (7) A total of 302,730 probes (targeting 300,341 sites, 32.52% of those on the array) are located in an enhancer or promoter, as indicated by a “TRUE” value in the “In_GeneHancer” column of the reannotated manifest.

Using both GENCODEv49 and GeneHancer, 825,504 sites (89.39% of all sites on the EPICv2 array) map to at least one gene or regulatory feature. For both intragenic and regulatory annotations, if an EPICv2 site is located within multiple overlapping transcripts, our annotation lists all transcripts. In total we provide an additional 19 columns of gene feature annotation for all sites on the EPICv2 array (Table 1). Figure 2 shows an example site in the manifest with its updated annotation and a guide to interpreting this information.

**Figure 2.**
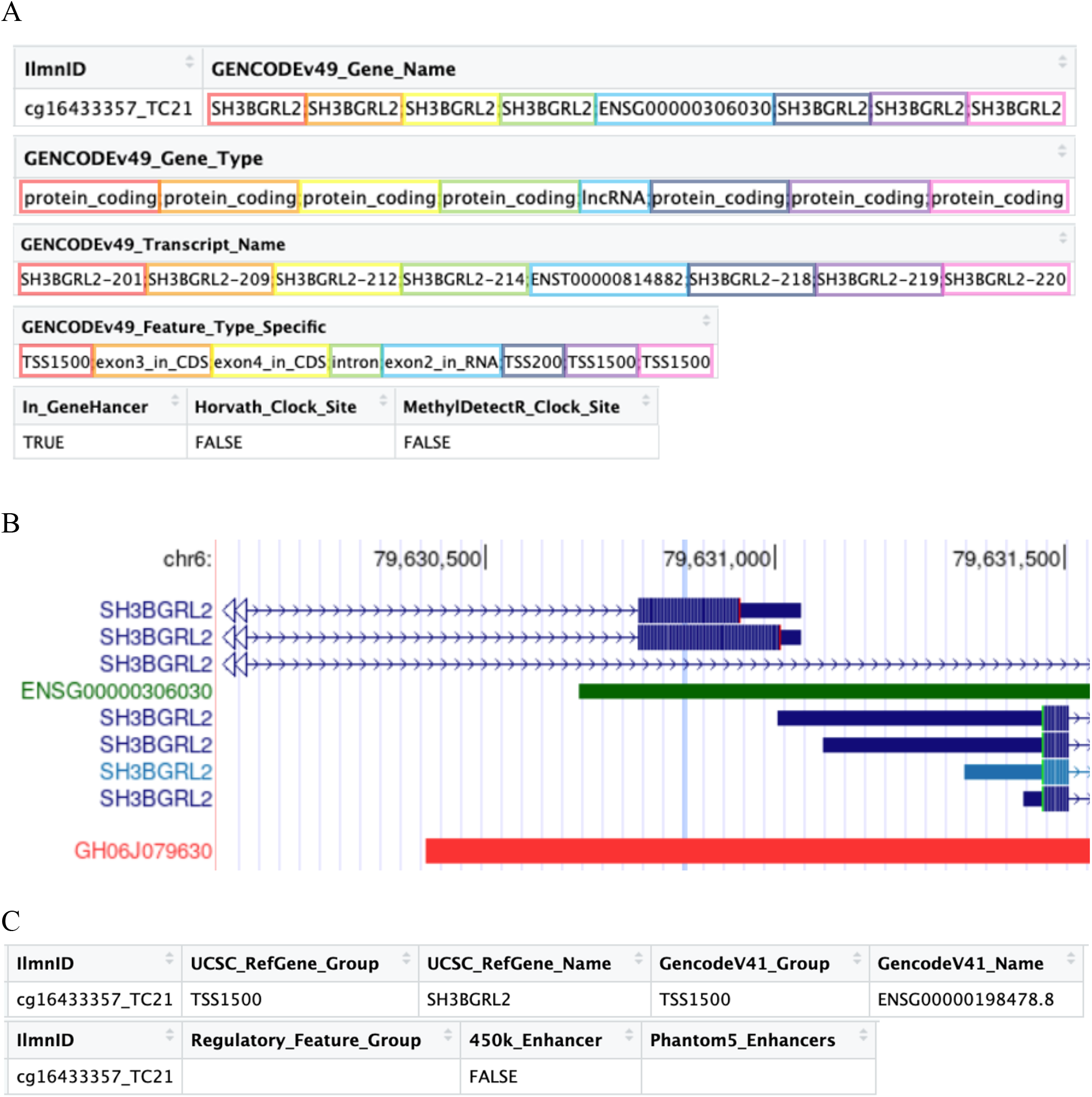
**A:** Screenshot of the re-annotated manifest, showing example annotation of site cg16433357. Only 7 of 19 new columns added to the manifest are shown for easier interpretation. As highlighted in the light blue boxes, this CpG site is located in exon 2 of one transcript of long non-coding RNA ENSG00000306030. It is also located in three different overlapping transcripts of the protein-coding gene SH3BGRL2, as shown by the orange, yellow and green boxes. For two of these transcripts (SH3BGRL2-209 and SH3BGRL2-212, highlighted in orange and yellow), the site is located in the coding region of exons 3 and 4 respectively, whereas the site is in an intron of the third transcript (SH3BGRL2-214, highlighted in green). The site is additionally located upstream of four other transcripts of SH3BGRL2 – <200bp 5’ of the TSS of transcript SH3BGRL2-218 (highlighted in dark blue), and 201-1500bp 5’ of transcripts SH3BGRL2-201, SH3BGRL2-219, and SH3BGRL2-220 (highlighted in red, purple, and pink). The “In_GeneHancer” column shows this site is within at least one promoter or enhancer in the GeneHancer database. The last two columns indicate that data from this site is not required as input to either the Horvath or MethylDetectR online uploaders. **B:** The region 500bp either side of site cg16433357_TC21 viewed with the UCSC Genome Browser. This confirms that the site overlaps three transcripts of gene SH3BGRL2 – being in located in an exon of two of the transcripts and the intron of another. The site is located <200bp upstream of the TSS of a fourth transcript of SH3BGRL2, and <1500bp upstream of three others. It is also located within exon 2 of RNA ENSG00000306030. Our annotation labelled this site as being within a GeneHancer regulatory element. Viewing the region with UCSC Genome Browser clarifies that this is element GH06J079630, a promoter that regulates genes including ELOVL4, LCA5, and SH3BGRL2. (21) **C:** Original annotation of site cg16433357_TC21 in the Illumina EPICv2 manifest. Illumina’s annotation correctly indicates that this site is within the TSS1500 of one transcript of SH3BGRL2 (ENSG00000198478 is the ENSEMBL ID of this gene (22)), but omits firstly that the site is promoter-associated, and secondly that the site is located in or <=1500bp upstream of six additional protein-coding transcripts of SH3BGRL2 and one transcript of ENSG00000306030.

### Epigenetic clock annotation

The Horvath DNA Methylation Age Calculator uses DNA methylation-based biomarkers of aging to predict the epigenetic age of samples, and is applicable to multiple cell/tissue types. (8) The original Horvath epigenetic clock used 353 loci to predict epigenetic age (8), but the more recent online uploader, which outputs the predictions of multiple epigenetic clocks, uses 94,394 loci. (9,17) The Horvath uploader technically allows for the upload of EPICv2 data – however, only 82,656 of the 94,394 sites are included on the EPICv2 array. We have added a column to the novel manifest – “Horvath_Multiple_Clock_Site” – labelling these sites with “TRUE” and all other EPICv2 sites as “FALSE”, allowing methylation data to be more easily filtered for upload to the Horvath clock. The 82,656 sites are captured by 83,548 probes, meaning there are 892 replicate probes, which the user will need to account for in their data prior to upload.

The MethylDetectR epigenetic clock outputs predictions of epigenetic age based on blood tissue, as well as predictions of multiple metabolic measures including BMI, alcohol consumption, smoking behaviour, etc. (10) Similar to the Horvath uploader, input to the MethylDetectR uploader is a matrix DNA methylation data (e.g. beta values) at 15,190 sites (11,18), of which 13,780 are included in the EPICv2 array. These have been labelled with “TRUE” in the “MethylDetectR_Clock_Site” column. 210 sites are read by replicate probes which will need to be de-duplicated.

### Comparison with Illumina’s EPICv2 manifest

The Illumina EPICv2 manifest annotates 542,165 sites (or 58.71% of all sites targeted by the EPICv2 array) as being located in any gene or regulatory feature – this includes sites mapped to exons, UTRs, within 1500bp upstream of the TSS, promoters, and enhancers, but not introns. In our version of the manifest, we annotate the aforementioned features as well as introns. This difference in included features, and our use of a more recent version of GENCODE, mean that we annotate 825,504 sites (89.39% of all sites on the EPICv2 array) to at least one gene or regulatory feature. If we exclude probes only annotated to introns, this reduces to 534,475 sites (57.88% of those on the array).

527,479 sites are mapped to a gene or regulatory feature in both our version of the manifest and the manufacturer’s. We annotate 298,025 sites to at least one gene or feature, which were not mapped to any in the manufacturer’s version of the manifest. 14,686 sites are no longer mapped to a gene or feature in our annotation (Figure 3A). Figure 3B shows that our annotation maps more sites to intragenic features, including exons, introns, and UTRs. However, we annotate less sites to the TSS200 or TSS1500.

**Figure 3.**
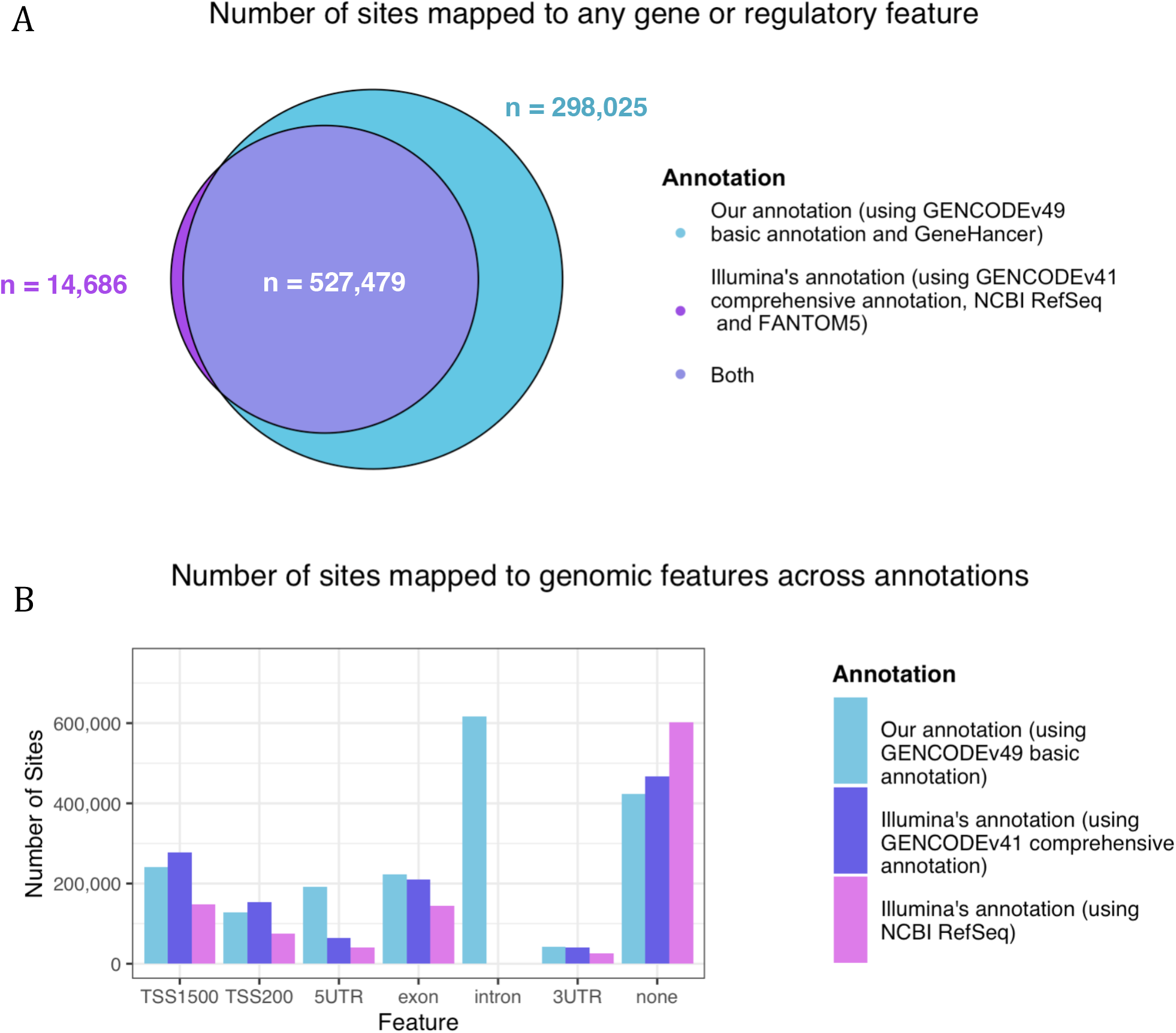
**A:** Venn diagram showing numbers of sites mapped to any gene or regulatory feature in our annotation only, the manufacturer’s annotation only, or both. **B:** Numbers of sites mapped to different genomic features according to our updated manifest (based on GENCODEv49 basic gene annotation) and the Illumina manifest (based on GENCODEv41 comprehensive annotation or NCBI RefSeq). Where probes are annotated to multiple different categories they are counted for each category.

### Updated manifest extends gene feature annotation to more than half of probes on EPICv2

214,808 sites were mapped to a gene body by the manufacturer. This includes all sites mapped to a coding or UTR exon. Comparatively, our version of the manifest maps 731,759 sites to a gene, and 222,963 sites to any exon. In total, 521,166 sites are newly labelled as intragenic in our annotation. This is 56.44% of all 923,452 genomic sites assayed by the array. 97.06% (505,860) of these sites are located in the intron of at least one transcript.

Of note, 4,215 sites were annotated as being within a gene body by the manufacturer, but are not in a gene body according to our annotation. 3,472 of these sites (82.37%) were labelled as being in a gene body according to the “GencodeV41_Group” column, whereas 1,182 of the sites (28.04%) were in a gene body according to the “UCSC_RefGene_Group” column. The manufacturer used the GENCODEv41 comprehensive gene set, whereas we used GENCODEv49 basic, which prioritises full-length, protein-coding transcripts in each gene. (19) Therefore, the sites which are no longer mapped to a gene in our annotation mostly reflect differences between these versions of GENCODE, but also some differences between GENCODEv49 and NCBI RefSeq, used by the manufacturer for the “UCSC_RefGene” annotation. (4)

### Differences in GENCODE reference sets reduce TSS-proximal annotations

The manufacturer’s version of the manifest maps 156,540 sites to at least one TSS200 region, whereas we annotate 128,789 sites as being in a TSS200. 110,466 sites are mapped to a TSS200 in both versions, whereas 18,323 sites are newly mapped to a TSS200 in our version, and 46,074 sites are no longer mapped to a TSS200.

281,255 sites were annotated as being in a TSS1500 by the manufacturer. We map less sites, 241,715, to a TSS1500. 213,302 sites are labelled as being in a TSS1500 in both versions, 28,413 are novel, and 67,953 are no longer mapped to a TSS1500.

Of the 105,869 total sites which are no longer mapped within 1500bp upstream of a TSS in our version of the manifest, 101,820 (96.17%) were labelled as being in a TSS200 or TSS1500 in the “GencodeV41_Group” column of the original manifest, whereas 10,492 sites (9.91%) were in a TSS200 or TSS1500 according to the “UCSC_RefGene_Group” column. In contrast to the GENCODEv41 comprehensive gene set, GENCODEv49 basic excludes numerous shorter transcripts with a TSS downstream of the canonical TSS, which reduces the number of sites annotated to a TSS200 or TSS1500 region – Supplementary Figure 1.

### Annotation with GeneHancer substantially increases the number of sites mapped to promoters and enhancers

Illumina label 113,342 sites as promoter-associated in the “Regulatory_Feature_Group” column of the original manifest. This includes both cell type-specific and general promoters. (4) Illumina also annotate 242,240 sites as within an enhancer, either according to the FANTOM5 database (20), or based on GENCODE-derived enhancer annotation for a previous version of the Infinium methylation array, the HumanMethylation450 BeadChip (450k array). (4) Using the GeneHancer database (which integrates FANTOM5), we annotated 300,341 sites as located in any enhancer or promoter. 152,567 of these sites (50.80%) were also in a regulatory element according to the manufacturer’s annotation, whereas 147,774 (49.20%) are newly annotated to a regulatory element in our version of the manifest. 19,489 (17.19%) of the sites annotated to a promoter by the manufacturer and 89,673 (37.02%) of the sites annotated to an enhancer are not annotated to any regulatory element in our annotation.

## Discussion

A comprehensive map of the genes and features profiled by the EPICv2 array is vital for interpreting the results of studies conducted using EPICv2. Our re-annotated manifest significantly increases the percentage of sites on the EPICv2 manifest mapped to any gene or regulatory feature, from 58.71% (542,165 sites) to 89.39% (825,504 sites). By improving coverage of the intragenic and regulatory genome, our annotation allows more complete assessment of the biological impact of differential methylation at sites of interest. Our annotation enables researchers to integrate findings and datasets from different versions of Illumina methylation BeadChips, by maintaining backwards compatibility with manifest files for previous iterations of the platform. The annotation file is available to researchers at https://doi.org/10.5281/zenodo.14933468.

Our version of the manifest particularly improves annotation of the non-coding genome, important because DNA methylation at promoters and enhancers regulates transcription, and DNA methylation at the exon-intron boundary impacts alternative splicing. (24,25) We used a subset of high-confidence regulatory elements from the GeneHancer database to annotate 32.52% of sites (300,341) as being in a promoter or enhancer. An advantage of this approach is that GeneHancer encompasses numerous databases, including the FANTOM5 database used by the manufacturer to label enhancers in the original manifest, widening the scope for promoter and enhancer discovery. However, one significant limitation is that we could not include detail on regulatory element names, or associated genes, due to GeneHancer data distribution restrictions. Instead we have indicated whether or not a site is present in a regulatory element, which will be sufficient for characterisation and enrichment analysis but limits functional interpretation. For this we recommend that users seek further information via UCSC Genome Browser and the GeneCards website.

The increased number of annotations in our updated manifest is primarily due to the inclusion of introns, which were excluded from the manufacturer’s annotation. Updating the original annotation from GENCODE release 41 to 49 also explains a portion of the difference, with 8.88% of sites (81,978) being newly annotated to an exon, TSS200, or TSS1500 region.

As gene annotation databases such as GENCODE and GeneHancer are frequently updated, further updates to the manifest may be required – our re-annotation code is therefore publicly available on GitHub. As our understanding of human genome structure, sequence variation, and functional annotation evolves, a transparent and consistent approach to re-annotation will be beneficial for epigenomic research.

There are some limitations which should be considered when using our manifest to interpret epigenetic analyses. Firstly, we solely annotated probes to genes and regulatory elements based on proximity. This strategy reflects the common assumption that DNA methylation preferentially influences the activity of local genes, promoters, and enhancers; however, it does not capture distal trans-regulatory effects. Secondly, we used the GENCODEv49 basic gene set, which prioritises high-confidence, full-length, protein-coding transcripts. Consequently, the re-annotated manifest may be less suitable for analyses focused on non-canonical and lower-confidence transcripts. Thirdly, our annotation includes isoforms from all genes. As transcript expression varies by tissue, cell type, and developmental stage, not all annotated transcripts will be relevant depending on biological context.

## Conclusion

We have re-annotated the EPICv2 manifest, using the latest versions of the GENCODE and GeneHancer public databases, providing an up-to-date and more comprehensive framework for interpreting DNA methylation data. The number of CpG sites assigned to genes has substantially increased in our annotation compared to the annotation provided by the manufacturer and maintains backwards compatibility with previous manifests. The re-annotation also identifies promoter and enhancer regions, and sites within the TSS200/1500. It offers a valuable resource for researchers, facilitating more accurate interpretation of the results of studies conducted using the EPICv2 array.

## Methods

The original EPICv2 manifest was downloaded from the Illumina website distributed as part of the Infinium MethylationEPIC v2.0 Product Files. (5) All analysis and re-annotation was conducted in the R programming language, version 4.3.2. (26)

### Intragenic/Gene Body Annotation

The GENCODE Human release 49 basic gene annotation, for genome version GRCh38.p14, was downloaded from the GENCODE website. (6,27) This annotation includes names, IDs, and coordinates of genes, transcripts, and exons for the whole human genome. The basic annotation prioritises high-confidence, protein-coding transcripts, and is described as “the main annotation file for most users” for release 42 onwards. (19,28,29)

When re-annotating the manifest, all EPICv2 sites located within a gene body were labelled with the IDs and names of any genes and transcripts they were located in, as well as whether the site was in an intron or exon, and the exon number if applicable. GENCODEv49 includes coordinates of features within exons (CDSs and UTRs) for protein coding genes only. Therefore, any EPICv2 sites within the exon of a protein coding gene were also annotated with this more granular level of detail, as being in a CDS or UTR.

The GENCODEv49 basic gene annotation does not include information about the location of introns. Therefore, we calculated intron coordinates for all genes as starting 1bp after the end coordinate of the preceding exon and 1bp before the start coordinate of the subsequent exon.

Sites profiled by the EPICv2 manifest were annotated as being within a feature (such as a gene, transcript, or intragenic feature including intron/exon/CDS/UTR) if 1. the genomic location of the site in genome build hg38 (found in the “MAPINFO” column of the manifest) fell on or between the start and end coordinates of the feature, and 2. the site and feature were located on the same chromosome. Figure 2 shows an example annotation for a probe, the target site of which is located in multiple overlapping transcripts. Table 1 describes annotation in more detail in the new version of the manifest.

### Distance-based Regulatory Feature Annotation

DNA methylation proximal to the TSS of a gene is more likely to affect expression levels of that gene – although this effect is primarily noted for the core promoter region (±50 base pairs (bp) around the TSS) (30) regulatory elements can span thousands of bp (31). For this reason, as well as annotating which EPICv2 array sites are located within a gene body, we also annotated which sites are located near the TSS.

To ensure our manifest is backwards-compatible with existing pipelines for EPICv2 data and earlier arrays, we replicated the approach in the manufacturer’s manifest of annotating sites in the TSS200 and TSS1500. We defined the TSS200 as the region 200bp 5’ of the TSS, i.e., if the genomic position of the TSS is N and the strand direction + then the TSS200 encompasses the region N-201 to N-1. If the strand direction is - then the TSS200 includes N+1 to N+201. The TSS1500 does not overlap the TSS200 in our annotation, encompassing N-1501 to N-202 on the + strand, and N+202 to N+1501 on the - strand. This is slightly different from Illumina’s annotation of the TSS200 and TSS1500, defined as the regions 1-200bp and 200-1500bp 5’ of the TSS respectively. (4)

We calculated the locations of TSS200 and TSS1500 regions using gene start coordinates accessed from GENCODEv49. We used the “MAPINFO” (genomic position) and “CHR” (chromosome) columns of the original manifest to determine which sites were located a TSS200/1500. These sites were labelled with “TSS200” and “TSS1500” respectively in the GENCODEv49_Feature_Type, GENCODE_v49_Feature_Type_Specific, and GENCODEv49_Feature_Exon_Number columns of the reannotated manifest – see Table 1 for description of all columns and contents.

### GeneHancer Regulatory Feature Annotation

GeneHancer is a database of regulatory features, part of the GeneCards database, which integrates multiple other databases to identify promoters/enhancers and predict their target genes. (7) GeneHancer double-elite annotation data was accessed via UCSC Table Browser. (14) The double-elite annotation includes only the regulatory elements where both the existence of the regulatory element, and its association with any linked genes, are supported by at least two sources of evidence. (7) It is not currently possible to download the entire GeneHancer database at once, however the user can download data for a specific region via UCSC Table Browser by specifying chromosome, start, and end coordinates of the region. GeneHancer table “GH Reg Elems (DE)” was downloaded for each chromosome from UCSC Table Browser by inputting chromosome start and end coordinates accessed via R package GenomeInfoDb version 1.42.3. (32)

Although the GeneHancer database includes the genomic coordinates and unique IDs of enhancers and promoters, as well as the genes which these elements regulate, data from GeneHancer cannot be redistributed in full or part without the permission of LifeMap Sciences. (13) Therefore, when re-annotating the manifest, we did not include annotation of which specific promoters/enhancers EPICv2 sites are located in, or linked genes. Instead we have included a binary column called “In_GeneHancer”, indicating whether or not a site is present within any promoter or enhancer according to the GeneHancer database. If a site of interest (for example, a site predicted to be significant following differential methylation analysis) is labelled “TRUE” in this column, this indicates that the site is located within a double-elite regulatory element. More information about the element and genes it regulates can be found online, e.g. by inputting the site’s name or genomic coordinates into UCSC Genome Browser (14) and viewing the GeneHancer track.

The “GH Reg Elems (DE)” table includes one row per regulatory element with columns “CHR”, “GeneHancer_Start”, and “GeneHancer_End” indicating chromosome and position of the element. Sites in the EPICv2 manifest were annotated as being within a regulatory element if the site was located on or between the start and end position of the element and on the same chromosome, determined using the “CHR” and “MAPINFO” columns of the original manifest. Based on this annotation, sites within any GeneHancer regulatory element were labelled with “TRUE” in the “In_GeneHancer” column of the re-annotated manifest, and all other sites labelled “FALSE”.

### Annotation of sites used in the Horvath DNA Methylation Age Calculator and MethylDetectR epigenetic clocks

EPICv2 sites required for upload to the Horvath DNA Methylation Age Calculator were identified by matching the “EPICv1_Loci” column in the original EPICv2 manifest, to the names of CpG sites in the file “datMiniAnnotation4_fixed.csv”, available to download from the Clock Foundation website. (17) Sites required for upload to MethylDetectR were identified by matching the “EPICv1_Loci” column to the CpGs listed in the “Truncate_to_these_CpGs.csv” file available from (18).

### Comparison of the original Illumina EPICv2 manifest and the re-annotated EPICv2 manifest

The EPICv2 array reads DNA methylation level at 923,452 genomic sites, using 937,690 probes. The EPICv2 manifest is a CSV file with one row per probe. (5) The number of unique sites targeted by the array was calculated by joining the “MAPINFO” column of the manifest (which gives the genomic location of the site being read by each probe) to the “CHR” column (the chromosome of the site), and counting the number of unique values, excluding any probes labelled as being on ‘chromosome 0’ or with NA values in these columns.

The original EPICv2 manifest uses GENCODEv41 (for columns “GencodeV41_Group”, “GencodeV41_Name”, and “GencodeV41_Accession”) and NCBI RefSeq data (for columns “UCSC_RefGene_Group”, “UCSC_RefGene_Name”, “UCSC_RefGene_Accession”) to annotate which sites are located in genes, transcripts, exons, UTRs or upstream of a TSS. (4,6,33) Importing GENCODEv41 comprehensive and basic annotation tracks into UCSC genome browser, then viewing a subset of randomly-chosen sites profiled by the EPICv2 array, revealed that the manufacturer most likely used the GENCODEv41 comprehensive annotation for all columns beginning “GencodeV41” (Supplementary Figure 2).

Any site labelled as being in a UTR or exon in either the “UCSC_RefGene_Group” or “GencodeV41_Group” columns of the original manifest was counted as being annotated to a gene body by the manufacturer. In the re-annotated manifest, we have removed the UCSC_RefGene and GencodeV41 annotations, replacing these with our annotation based on the GENCODEv49 basic gene set release. Using a grep search, any site containing the terms “exon”, “intron”, and/or “UTR” in the “GENCODEv49_Feature_Type” column was counted as being annotated to a gene body in the novel manifest.

The number of sites labelled as being in the TSS200 and TSS1500 by Illumina was counted by looking for the terms “TSS200” and “TSS1500” respectively in the “UCSC_RefGene_Group” and/or “GencodeV41_Group” columns. We compared this to the number of sites labelled with “TSS200” or “TSS1500” in the “GENCODEv49_Feature_Type” column.

Sites annotated as being in a promoter by Illumina were identified by searching for the term “Promoter” in the “Regulatory_Feature_Group” column of the original manifest. Sites annotated as being in an enhancer by Illumina were identified by looking for a value of “TRUE” in the “450k_Enhancer” column or any non-empty value in the “Phantom5_Enhancers” column. This was compared to the number of sites annotated with “TRUE” in the “In_GeneHancer” column of the re-annotated manifest.

## Contributions

BMR wrote the manuscript, re-annotated the EPICv2 manifest with genes/transcripts/intragenic features using GENCODEv49 data, added the GeneHancer annotation, added the annotation of which sites are required for upload to epigenetic clocks, formatted the manifest for readability and uploaded it publicly.

BMR and AW conceptualised and planned the project of re-annotating the manifest.

AW, EH, and JM proofread the manuscript and provided feedback and ideas.

AW and EH supervised the project.

Other acknowledgements

The authors thank Dr. Philippa Wells for extensive feedback on the project, and contributing ideas for future work. Thank you to Alice Franklin for proofreading the manuscript. Finally, we would like to thank the Complex Disease Epigenomics Group at the University of Exeter for their support and feedback.

