## Supplementary Figures for "Re-annotating the EPICv2 manifest with genes, intragenic features, and regulatory elements"

A

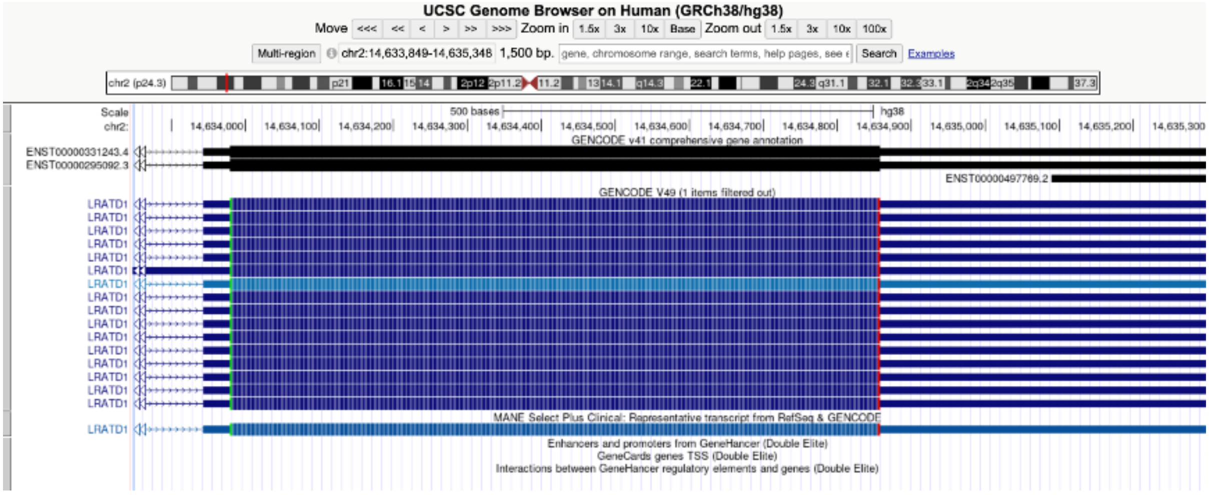


B

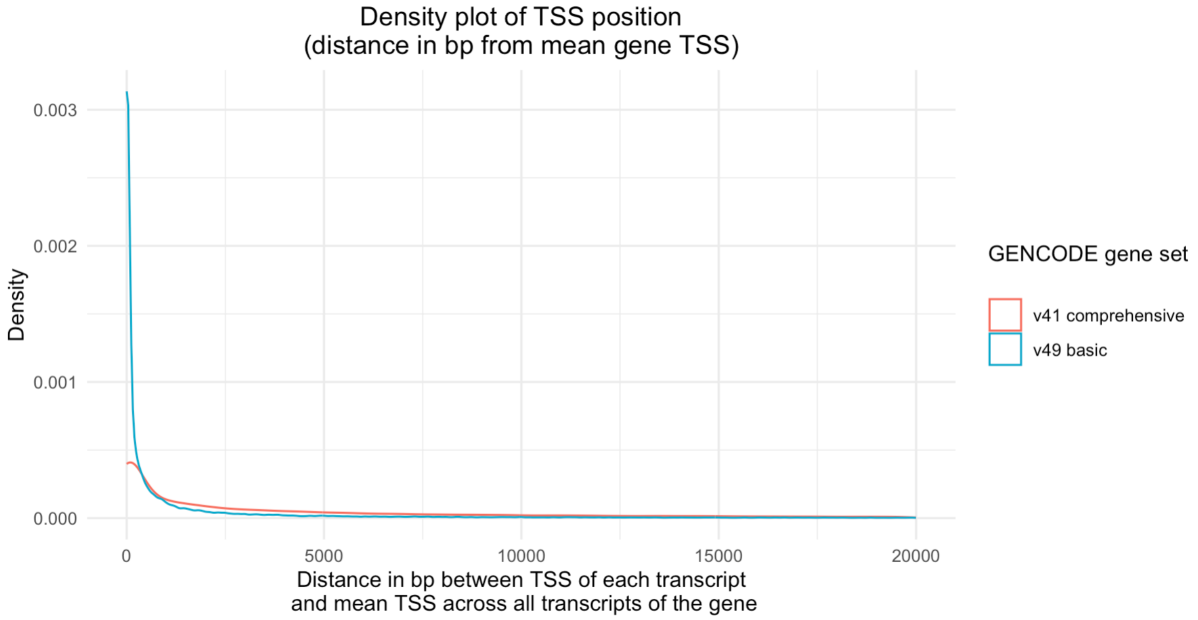


**Supplementary Figure 1A:** Example CpG site cg16846645 viewed in UCSC genome browser, with v41 comprehensive and v49 basic GENCODE tracks shown. The CpG site is highlighted in blue on the left side of the screenshot. Sites mapped to a TSS200/TSS1500 region in the manufacturer's annotation but not ours were selected at random to view. As in this example, discrepancies exist due to shorter transcripts such as ENST00000497769.2 being excluded from the GENCODE v49 basic gene set.

**B:** Density plot, including the majority of transcripts in GENCODE v41 comprehensive and v49 basic, of distance in bp between transcript TSS and mean TSS of its gene. 202,249/251,063 (80.56%) of transcripts from GENCODEv41 comprehensive and 259,955/280,000 (92.84%) of transcripts from GENCODEv49 basic were included. Transcripts with a TSS >20,000bp away from mean TSS for the gene were excluded for clearer visualisation. This shows that TSS position is much more conserved across transcripts of the same gene in the GENCODEv49 basic gene set relative to GENCODEv41 comprehensive. Increased variation in TSS position in the latter may explain why numerous sites have been annotated as being within 1500bp 5' of a TSS in the manufacturer's annotation, but not in ours.

A

| IlmnID | CHR | MAPINFO | GencodeV41_Group | GencodeV41_Name | GENCODEv49_Gene_Name | GENCODEv49_Feature_Type |
|---|---|---|---|---|---|---|
| cg03565437_TC11 | chr5 | 14488373 | exon_43;TSS200 | TRIO;ENSG00000038382.22 | TRIO | intron |

B

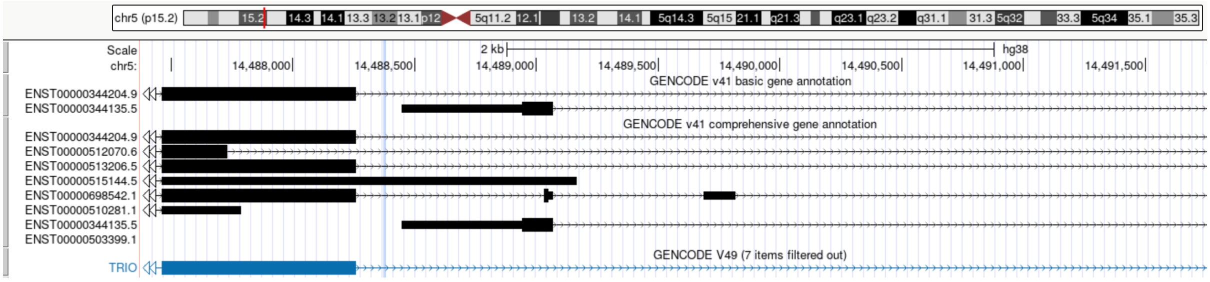


**Supplementary Figure 2A:** Screenshot of the re-annotated manifest, showing the manufacturer's annotation of site cg03565437 using GENCODEv41 and our annotation using GENCODEv49. The manufacturer's annotation indicates this site is in an exon of TRIO and <=200bp 5' of ENSG00000038382. Whereas our annotation labels it as being in an intron of TRIO only.

**B:** Importing the GENCODEv41 basic and comprehensive annotations as custom tracks into UCSC Genome Browser shows the difference in annotations is explained by both GENCODE version and differences in the comprehensive/basic annotation. Gene ENSG00000038382 (transcript ENST00000344135.5, in the screenshot) exists in both the basic and comprehensive GENCODEv41 sets, but not in GENCODEv49, hence its exclusion from our annotation.

The CpG site (highlighted in blue) is within the intron of one transcript of TRIO (ENST00000344204.9) in both the GENCODEv41 and v49 basic gene annotations. It is within an exon of one transcript of TRIO (ENST00000515144.5), in the comprehensive annotation only. Although descriptions of the EPICv2 manifest columns provided by the manufacturer do not indicate which GENCODEv41 annotation was used to create the manifest (4), examples such as this suggest it was the GENCODEv41 comprehensive gene annotation.
